# Phylogenetic analyses reveal bat communities in Northwestern Mexico harbour a high diversity of novel cryptic ectoparasite species

**DOI:** 10.1101/2022.08.09.503385

**Authors:** Laura A. Najera-Cortazar, Alex Keen, Thomas Kitching, Drew Stokes, Simon J. Goodman

## Abstract

**Background:** Parasites are integral parts of ecosystem function and important drivers of evolutionary processes. Characterising ectoparasite diversity is fundamental to studies of host-parasite interactions, evolution, and conservation, and also for understanding emerging disease threats for some vector borne pathogens. With more than 1,400 species, bats represent the second most speciose mammalian clade, but their ectoparasite fauna are poorly known for most species.

**Methods:** We sequenced mitochondrial Cytochrome Oxidase C subunit I and nuclear 18S ribosomal gene fragments, and used Bayesian phylogenetic analyses to characterise ectoparasite taxon identity and diversity for 17 species of parasitized bats sampled along the Baja California peninsula and in Northwestern Mexico.

**Results:** The sequence data revealed multiple novel lineages of bat bugs (Cimicidae), flies (Nycteribiidae and Streblidae) and ticks (Argasidae). Within families, the new linages showed more than 10% sequence divergence, which is consistent with separation at least at the species level. Both families of bat flies showed host specificity, particularly on *Myotis* species. We also identified new records for the Baja peninsula of one tick (*Carios kelleyi*), and of five Streblid bat flies. One Nycteribiid bat fly haplotype from Pallid bat (*Antrozous pallidus*) hosts was found throughout the peninsula, suggesting potential long distance co-dispersal with hosts. Different bat bug and tick communities were found in the north and south of the peninsula.

**Conclusions:** This study is the first systematic survey of bat ectoparasites in the Baja California peninsula, discovering highly genetically differentiated novel lineages compared to other parts of North America. For some ectoparasite species, haplotype distributions may reflect patterns of bat migration. This work is a first step in characterising ectoparasite diversity over the Baja California peninsula, and understanding how ecological and evolutionary interactions shape bat ectoparasite communities among host species in different parts of their ranges.

## Introduction

Characterising ectoparasite diversity is fundamental to studies of host-parasite interactions, parasite evolution and conservation, and for understanding emerging disease threats for some vector borne pathogens (Morand, Krasnov, & Poulin, 2006; Poulin, 2014; Spencer & Zuk, 2016). Bat ectoparasites are of particular interest as model systems for host-parasite co-evolution and parasite community structure studies (Gómez & Nichols, 2013; Spencer & Zuk, 2016). Bat-ectoparasite relationships are also important for understanding bat dispersal, pathogen transmission and zoonotic disease risks (Klimpel & Mehlhorn, 2014; Speer et al., 2019; Wilder, Kunz, & Sorenson, 2015). Despite the widely recognised need to increase sampling effort for pathogen/parasite discovery in bats, bat ectoparasites are still understudied in most parts of the world (Gay et al., 2014; Reinhardt & Siva-Jothy, 2007). Bat-associated ectoparasites include Insecta, comprising bat bugs (Hemiptera), fleas (Siphonaptera) and flies (Diptera); and Arachnida comprising ticks (Ixodida) and mites (orders Mesostigmata and Trombidiformes) (Seneviratne, Fernando H, & Udagama-Randeniya, 2009; Ter Hofstede & Fenton, 2005). All these ectoparasites are hematophagous (blood feeding) organisms, creating potential zoonotic disease transmission risks for humans and domestic animals (Poulin, 2014).

Bat bugs (family Cimicidae), have a worldwide distribution and comprise approximately 110 known species within 24 genera, the majority of which are ecologically and biologically associated with bats (Hornok et al., 2017; Ossa et al., 2019). A second, less studied family, Polyctenidae also includes bat-associated bugs, with tropical and subtropical distributions (Klimpel & Mehlhorn, 2014).

Bat flies comprise two main families, Nycteribiidae and Streblidae, which have a common origin from a single lineage coevolving with bats (Dittmar, Porter, Murray, & Whiting, 2006; Wenzel & Tipton, 1966). As of 2006, there were approximately 520 species described (Dittmar et al., 2006). While new records are added regularly (Saldaña-Vázquez, Sandoval-Ruiz, Veloz-Maldonado, Durán, & Ramírez-Martínez, 2019; Szentiványi, Christe, & Glaizot, 2019), taxonomy at the species level is often ambiguous, without an updated review yet to be published. Additional fly families associated with bats include the Hippoboscidae and Glossinidae, which also parasitize other mammals and birds (Petersen, Meier, Kutty, & Wiegmann, 2007).

Ticks are grouped in three families: Argasidae (soft ticks), Ixodidae (hard ticks) and Nuttalliellidae. The latter family comprises a single known species, *Nuttalliella namaqua* (Burger, Shao, Labruna, & Barker, 2014; Mans, de Klerk, Pienaar, de Castro, & Latif, 2012), which has been suggested as the basal tick lineage (Mans et al., 2012). There are approximately 894 known species of ticks worldwide, with 32 species of Argasidae and 68 species of Ixodidae in Mexico (Guglielmone et al., 2010; Pérez, Guzmán-Cornejo, Montiel-Parra, Paredes-León, & Rivas, 2014). Bat tick phylogenetic studies have mainly focused on old world species (Hornok et al., 2017), with a limited number for the Americas and other parts of the world (Black IV, Klompen, & Keirans, 1997; Burger et al., 2014), and suggest bat-associated ticks commonly exhibit host-specificity (Sándor et al., 2019). Classification of taxonomic relationships among soft ticks are controversial (Burger et al., 2014), and more studies are needed to accurately describe their taxonomic status (Burger et al., 2014; Estrada-Pena et al., 2010).

Previous bat ectoparasite studies in North America have described the distribution and taxonomic status of bat flies, bugs and ticks (Bradshaw & Ross, 1961; Graciolli, Autino, & Claps, 2007; Jobling, 1938; Usinger, 1966), and their medical importance (Dick, Gannon, Little, & Patrick, 2003; Gill, Rowley, Bush, Viner, & Gilchrist, 2004; Steinlein, Durden, & Cannon, 2001), from a limited number of bat host species. In Mexico, publications have focused on describing species diversity of flies, ticks and mites (Bolaños-García, Rodríguez-Estrella, & Guzmán-Cornejo, 2018; Guzmán-Cornejo et al., 2017; Pérez et al., 2014); new records of bat flies (Cuxim-Koyoc, Reyes-Novelo, Morales-Malacara, Bolívar-Cimé, & Laborde, 2015; Ramírez-Martínez et al., 2016; Trujillo-Pahua & Ibáñez-Bernal, 2018); ecological relationships between bats and bat flies (Saldaña-Vázquez et al., 2019; Salinas-Ramos, Zaldívar-Riverón, Rebollo-Hernández, & Herrera-M, 2018; Zamora-Mejías, Morales-Malacara, Rodríguez-Herrera, Ojeda, & Medellín, 2020); and *Rickettsia* presence in soft ticks in Mexico (Sánchez-Montes et al., 2016). To our knowledge, there have been no studies of bat bugs in Mexico.

Baja California (Figure 1) is an isolated peninsula, 1300km long, in northwestern Mexico. Its complex geological history means it has a mosaic of habitats, with high levels of biodiversity and endemism (González-Abraham, Garcillán, & Ezcurra, 2010; Nájera-Cortazar, Álvarez-Castañeda, & De Luna, 2015; Riddle, Hafner, Alexander, & Jaeger, 2000), with approximately 25 species of bats recorded (Álvarez-Castañeda, Álvarez, & González-Ruíz, 2015; Medellín, Arita, & Sánchez, 2007). While there are several studies describing Baja bat diversity, distributions and ecology (Álvarez-Castañeda et al., 2015; Frick, Hayes, & Heady, 2008; Nájera-Cortazar et al., 2015), there are no studies describing the bat ectoparasite fauna.

**Figure 1.**
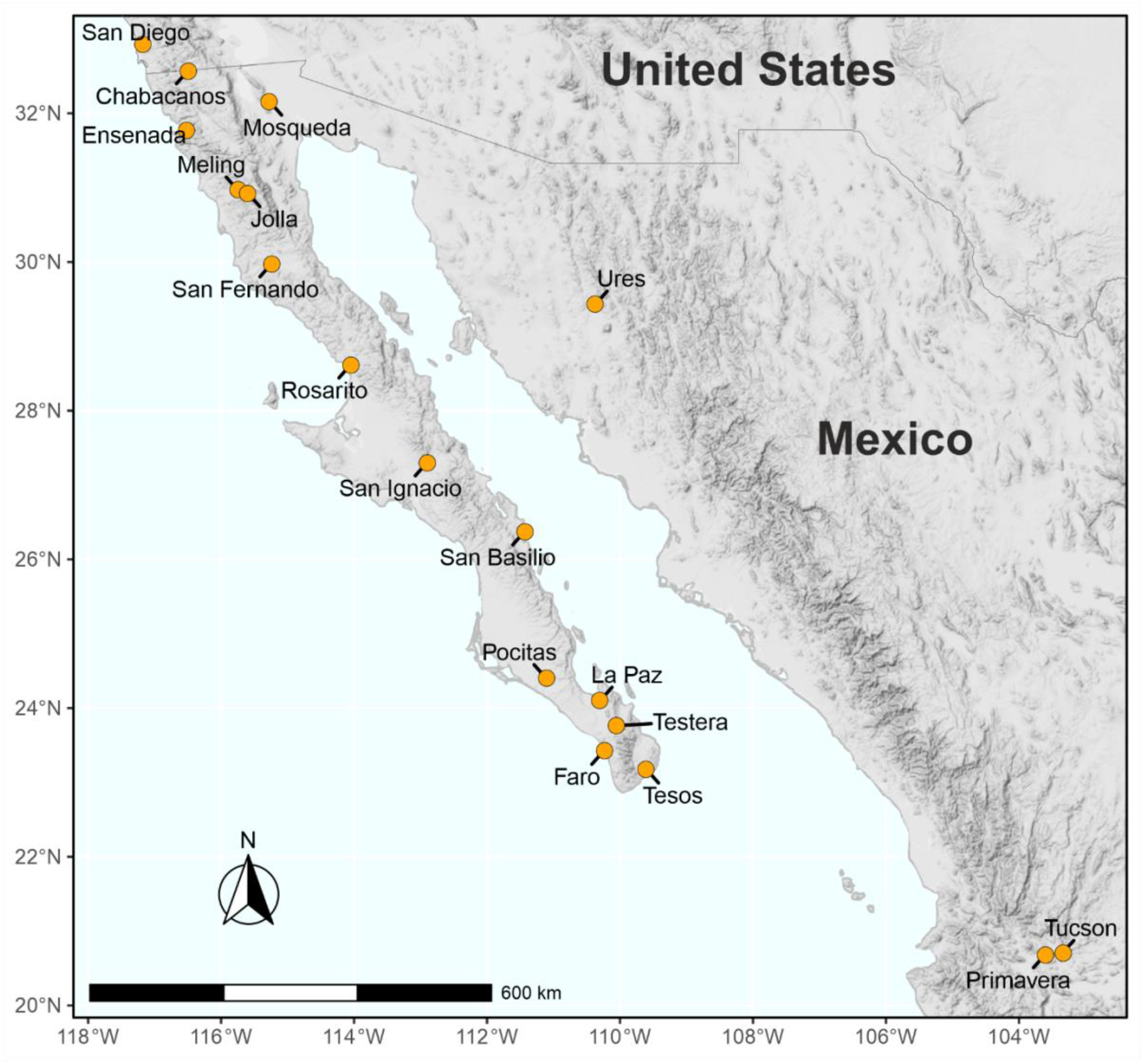
Map of sampling locations for bats and bat ectoparasites reported in this study.

Here we use a phylogenetic approach to characterise ectoparasite (bat bugs, flies, and ticks) identity and diversity for17 species of bat resident along the Baja California peninsula, and northwestern Mexico, based on mitochondrial Cytochrome Oxidase C subunit I (COI) and nuclear 18S ribosomal (18S) DNA amplicon sequences. Ectoparasites are assigned as known or novel lineages within Baja and the rest of the Americas, and we evaluate potential distributions with respect to Baja, and their hosts’ wider ranges. The results are relevant for increasing knowledge of bat and ectoparasite distributions across western North America, and to provide a basis for understanding how ecological and evolutionary interactions shape parasite community structure along environmental gradients such as those found in northwestern Mexico.

## Methods

### Sample collection

Bat sampling and collection of ectoparasites was conducted at 14 sites along the Baja California peninsula, and at three sites in western continental Mexico during 2016-2018 (Figure 1). Bats were identified to species level in the field following published identification guides (Álvarez-Castañeda et al., 2015; Medellín et al., 2007), and subsequently, *Myotis* bats were identified by molecular assays (Najera-Cortazar, 2020). A list of sites and bat species sampled is given in Table 1. Ectoparasites were collected manually from bats using forceps and transferred to labelled Eppendorf tubes with 96% ethanol for storage. Ectoparasites collected from the same bat but from different taxonomic families were stored in separate tubes for each family, with corresponding labels and appropriate specimen source identifiers. All samples were kept on ice during fieldwork, until their arrival to the laboratory where they were stored at −20° C. Ectoparasites were photographed during fieldwork using a portable Maozua 5MP 20-300X USB microscope, taking ventral and dorsal views. Preliminary classifications of ectoparasites were assigned according to morphological characters specified in keys by McDaniel (McDaniel, 1979), Usinger (Usinger, 1966), Knee and Proctor (Knee & Proctor, 2006) and Dick and Miller (Dick & Miller, 2010), and adapted for North American ectoparasites by this study. In addition, 10 specimens of bat bugs from *Parastrellus hesperus* hosts, captured at Ramona, California, U.S.A. (Site 0), were donated from the San Diego Natural History Museum, California U.S.A. Representative photographs for individuals of each ectoparasite genetic lineages identified in the study, without sexual determination, are shown in appendices (Appendices A1-A3).

**Table 1.**
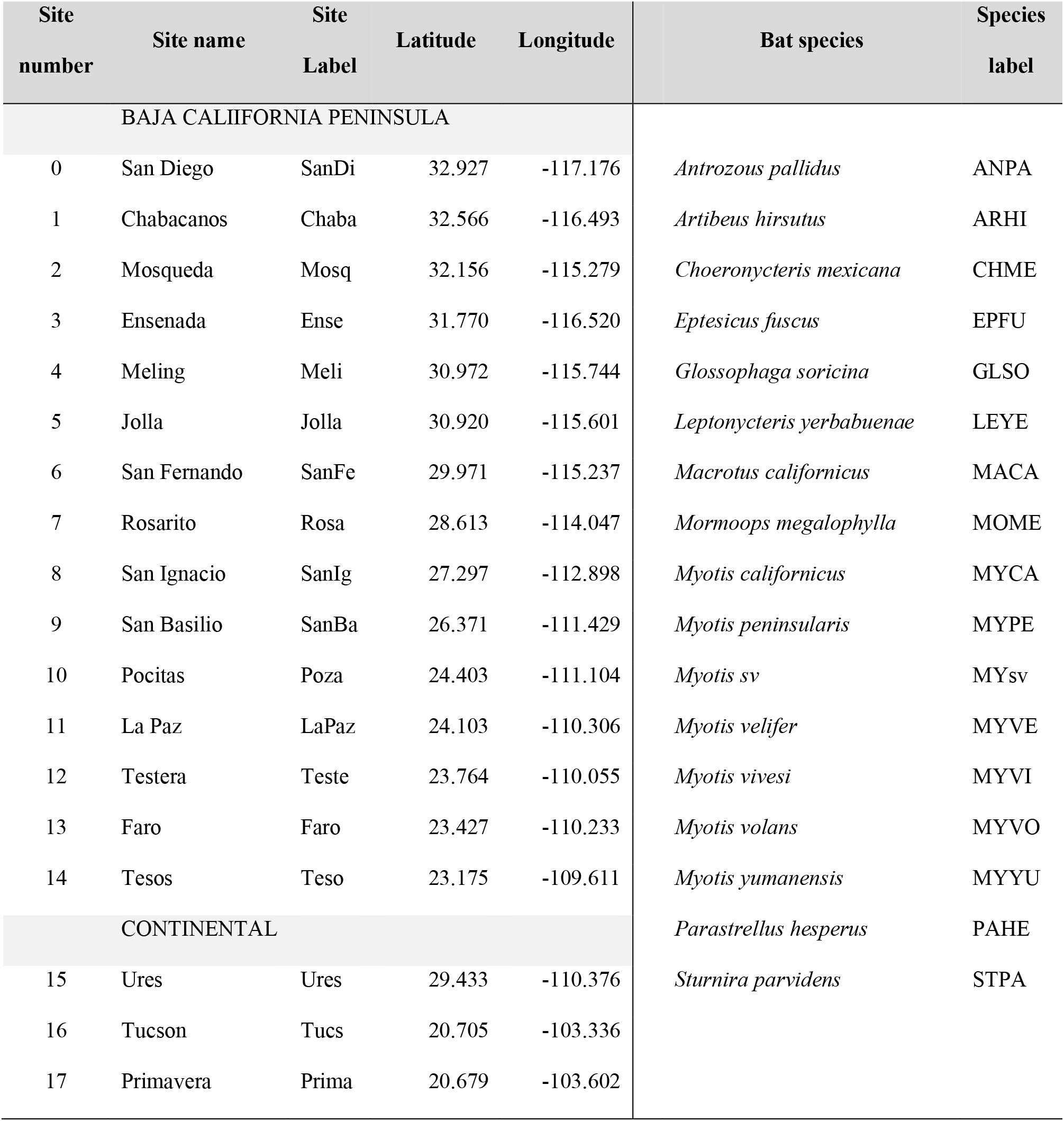
Left section: List of sites sampled showing number, name, abbreviation (Site label) and geographic coordinates. Right section: List of bat species sampled and their short labels (Species label). *Myotis sv* refers to any unidentified *Myotis* morphologically resembling *M. velifer*, found at Ures site.

| Site<br>number | Site name | Site<br>Label | Latitude | Longitude | Bat species | Species<br>label |
| --- | --- | --- | --- | --- | --- | --- |
| BAJA CALIIFORNIA PENINSULA |  |  |  |  |  |  |
| 0 | San Diego | SanDi | 32.927 | -117.176 | *Antrozous pallidus* | ANPA |
| 1 | Chabacanos | Chaba | 32.566 | -116.493 | *Artibeus hirsutus* | ARHI |
| 2 | Mosqueda | Mosq | 32.156 | -115.279 | *Choeronycteris mexicana* | CHME |
| 3 | Ensenada | Ense | 31.770 | -116.520 | *Eptesicus fuscus* | EPFU |
| 4 | Meling | Meli | 30.972 | -115.744 | *Glossophaga soricina* | GLSO |
| 5 | Jolla | Jolla | 30.920 | -115.601 | *Leptonycteris yerbabuenae* | LEYE |
| 6 | San Fernando | SanFe | 29.971 | -115.237 | *Macrotus californicus* | MACA |
| 7 | Rosarito | Rosa | 28.613 | -114.047 | *Mormoops megalophylla* | MOME |
| 8 | San Ignacio | SanIg | 27.297 | -112.898 | *Myotis californicus* | MYCA |
| 9 | San Basilio | SanBa | 26.371 | -111.429 | *Myotis peninsularis* | MYPE |
| 10 | Pocitas | Poza | 24.403 | -111.104 | *Myotis sv* | MYsv |
| 11 | La Paz | LaPaz | 24.103 | -110.306 | *Myotis velifer* | MYVE |
| 12 | Testera | Teste | 23.764 | -110.055 | *Myotis vivesi* | MYVI |
| 13 | Faro | Faro | 23.427 | -110.233 | *Myotis volans* | MYVO |
| 14 | Tesos | Teso | 23.175 | -109.611 | *Myotis yumanensis* | MYYU |
| CONTINENTAL |  |  |  |  | *Parastrellus hesperus* | PAHE |
| 15 | Ures | Ures | 29.433 | -110.376 | *Sturnira parvidens* | STPA |
| 16 | Tucson | Tucs | 20.705 | -103.336 |  |  |
| 17 | Primavera | Prima | 20.679 | -103.602 |  |  |

### DNA extraction, primer selection and PCR amplification

Individual ectoparasite bodies were crushed in Eppendorf tubes using sterile pestles, followed by digestion with Proteinase *K* at 56° C for 18 hours. DNA was extracted using either a Thermo-Fisher DNA extraction and purification kit (Thermo Fisher Scientific Inc.) or QuickExtract kit (Epicentre, Illumina), following the manufacturers’ protocols. PCR conditions were optimized separately for each ectoparasite group, and according to each primer set. Primers LCO1490 (5’-GGTCAACAAATCATAAAGATATTGG-3) and HCO2198 (5’-TAAACTTCAGGGTGACCAAAAAATCA-3’) (Folmer, Black, Hoeh, Lutz, & Vrijenhoek, 1994) were used to amplify an approximately 700bp fragment of the mitochondrial COI gene in a 25 μl reaction containing: 1 U of Flexi GoTaq Taq Polymerase, 5X GoTaq reaction buffer, 50 mM MgCl_2_ (1.5 – 2.5 mM final concentration), 0.5 μl PCR nucleotide Mix (0.2 mM each), 0.5 μl of each set of primers (1 μM final concentration), 15.8 μl ddH_2_O and 8 μl template DNA (extractions diluted at 1:10 with sterile distilled water). Thermal cycling parameters were as follows on a TECHNE thermocycler model TC-512: initial denaturation step at 95°C for 5 min, followed by 40 cycles of denaturation at 94°C for 40 seconds, annealing at 53°C - 56°C for 1 minute and extension at 72°C for 1 minute. Final extension was performed at 72°C for 10 minutes.

For the 18S rDNA gene, approximately 800bp amplicons were amplified using the primers 18S-1F (5’ CTGGTTGATCCTGCCAGTAGT 3’) and 18S-3R (5’ GGTTAGAACTAGGGCGGTATCT 3’) for bat bugs (Campbell, Steffen-Campbell, Sorensen, & Gill, 1995); a0.7-F (5’ ATTAAAGTTGTTGCGGTT 3’) and 7R (5’ GCATCACAGACCTGTTATTGC 3’) for flies (Whiting, 2002); and D-F (5’ GGCCCCGTAATTGGAATGAGTA 3’) and C-R (5’ CTGAGATCCAACTACGAGCTT 3’) for ticks (Mangold, Bargues, & Mas-Coma, 1997). The same reaction mix quantities were used as for the COI gene, with MgCl_2_ at a final concentration of 1.5 mM for all primer sets. Thermal cycling parameters were: initial denaturation at 95°C for 5 minutes, followed by 30 cycles of denaturation at 94°C for 40s, annealing at 53°C for 1 min and extension at 72°C for 1 minute. Final extension was performed at 72°C for 10 minutes.

PCR products were visualized by gel electrophoresis using 1% agarose with GelRed® (Biotium Ltd) staining, and then sent for PCR product purification and Sanger sequencing to Genewiz Inc. (Azenta Life Sciences, Leipzig, Germany), with each amplicon sequenced in both forward and reverse directions.

### Quality control and references sequences

Sequence quality for both the forward and reverse strands of each amplicon was evaluated in BioEdit 7.2.5 (Hall, 2005). Trimmed forward and reverse sequences were combined to generate a consensus sequence for each amplicon, and then analysed in BLAST (The National Library of Medicine, 2018) to generate initial taxon identities and identify reference sequences. Additionally, COI sequences were also analysed in the BOLD system platform (Ratnasingham & Hebert, 2007). Reference sequences for phylogenetic analyses were compiled from previous studies on each ectoparasite group (Burger et al., 2014; Dittmar et al., 2006; Mans et al., 2012; Tortosa et al., 2013), and by performing a systematic search using *AnnotationBustR* 1.2 package (Borstein, 2018) in RStudio 1.1.456 (RStudio Team, 2021), searching for the closest genus for the sequences generated in this study and for those obtained by BLAST. Alignments combining reference sequences and those from this study were generated using CLUSTAL W (Thompson, Higgins, & Gibson, 1994), implemented BioEdit, and were reviewed by eye, with manual correction of potential errors as required.

### Sequence summary statistics and phylogenetic analyses

Genetic distance estimates among haplotypes and best fit sequence evolution models for the COI and 18S datasets were evaluated using MEGA 10.1.7 (Sudhir, Stecher, Li, Knyaz, & Tamura, 2018). The Barcode Index Number system was followed to delimit a lineage, where intraspecific variation at COI is generally considered as groups of sequences with less than 2% divergence, and exhibiting more than 4% divergence from neighbouring lineages (Hebert, Ratnasingham, & DeWaard, 2003; Ratnasingham & Hebert, 2013; Salinas-Ramos et al., 2018).

Bayesian phylogenetic analyses were performed using the best fit evolution model identified for each group and each marker implemented with BEAST 1.10.4 (Suchard et al., 2018). A MCMC chain length of 10,000,000 was used and priors specific to each parasite group and sequence evolution model selected using the program BEAUti 1.10.4 (Suchard et al., 2018). Phylogenetic analyses were performed separately for each ectoparasite group and each marker. Nycteribiidae and Streblidae families of bat flies were analysed separately in order to improve sequence alignment quality and phylogenetic resolution. For each family-marker combination, two separate runs using the same Bayesian settings file generated by BEAUti were run in BEAST, with a burn in of the 10% of the total number of iterations. After, stationarity of BEAST results were assessed in Tracer 1.7.1 (Rambaut *et al*., 2018). Both files for each ectoparasite family-marker combination were combined using LogCombiner 1.10.4 (Suchard *et al*., 2018), generating a single .log file and a single .tree file. A majority-rule consensus tree was inferred using TreeAnnotator 1.10.4 (Suchard *et al*., 2018) with the combined .tree file, and using a posterior probability limit of 0.6, median nodes heights were summarised. Phylogenetic trees were annotated, edited and visualized using iTOL 5.6.2 (Letunic & Bork, 2016).

Following phylogenetic analysis, haplotypes for each family were grouped by lineage, and haplotype diversity and genetic summary statistics were calculated in DNAsp 5.10.01 (Librado & Rozas, 2009). Outgroups were chosen based on previous phylogenetic studies of each group, using *Bucimex chilensis, Primicimex cavernis* and *Anthocoris flavipes* for bat bugs (Ossa et al., 2019)(Ossa et al., 2019)(Ossa et al., 2019)(Ossa et al., 2019); *Drosophila melanogaster, Chrysops niger, Musca domestica*, and *Sarcophaga bullata* (Dittmar et al., 2006), for both bat fly families, plus *Ornithomya avicularia* for the family Streblidae only; and *Nuttalliella namaqua* (Mans et al., 2012) for ticks.

## Results

### Sampling and field identifications

A total of 1,988 ectoparasites were collected within the orders Diptera (flies), Hemiptera (bugs), Ixodida (ticks), Mesostigmata (mites), Siphonaptera (fleas) and Trombidiformes (chiggers). Fleas, chiggers, mites and any unclassified specimens (e.g. where it was not possible to differentiate small tick larvae and mites) were excluded for the present study. The remaining samples comprised 90 bat bugs, 213 bat flies and 126 ticks, collected from 138 individual bat hosts (Figure 2) of 17 species.

**Figure 2.**
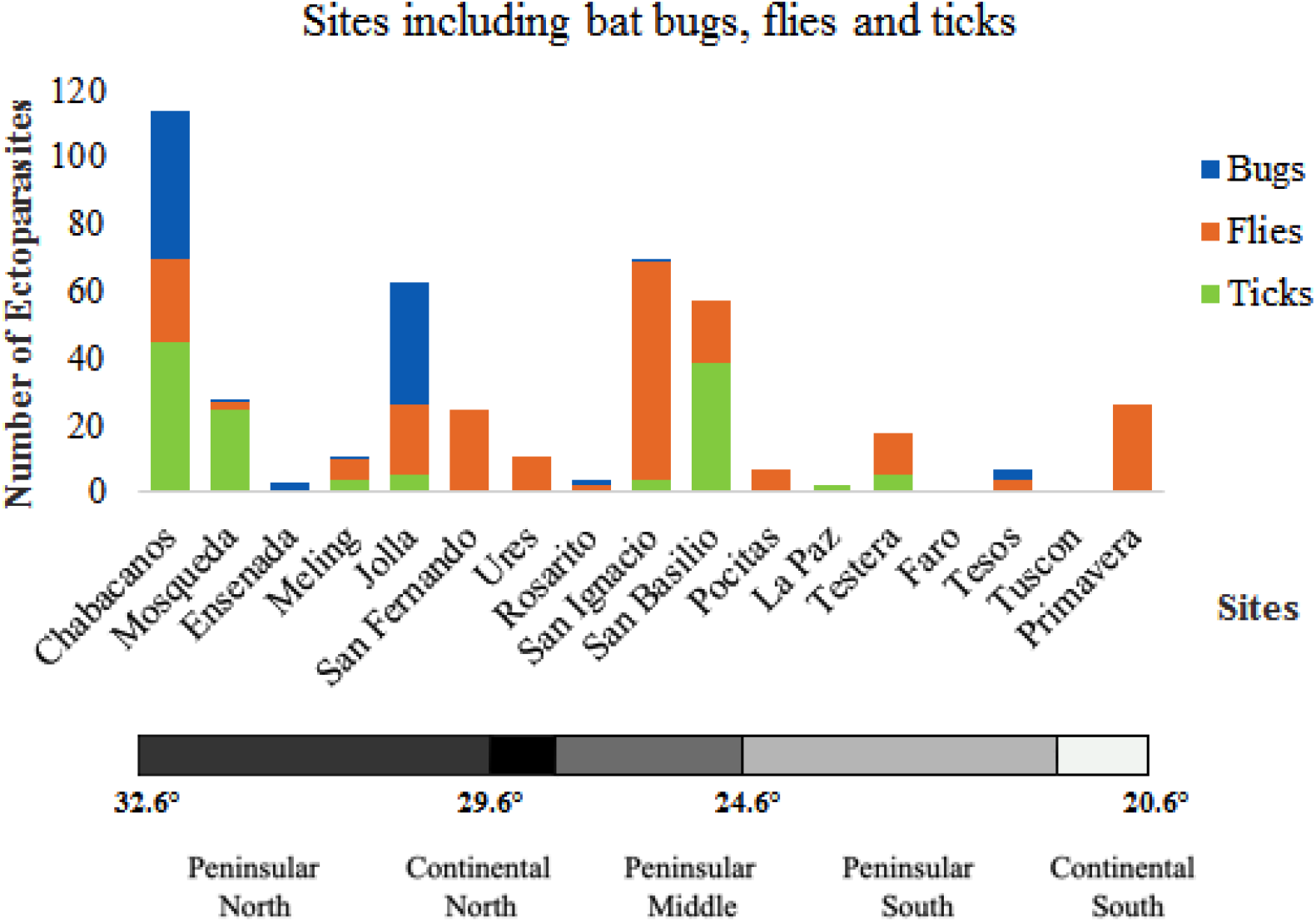
Number of bat bugs, flies and ticks captured per site in 2016-2018 fieldwork.

Flies were the predominant ectoparasite type, found in 13 of 17 sites, followed by ticks and bugs, present in eight sites each. Initial field morphological evaluations suggested most bug specimens were related to *Cimex pilosellus* (Usinger, 1966). Most bats flies could be identified to the genus level, which were later corroborated with molecular data. On the basis of morphology, all bat ticks were identified as family Argasidae (soft ticks), with at least six different morphotypes present. Most of these were tentatively attributed to the genus *Ornithodoros*.

### BLAST and BOLD sequence matches

Amplicon sequences were generated from 145 specimens for mitochondrial COI and from 147 specimens for 18S rDNA across the three groups: bugs *n* = 30/30 (COI/18S, respectively); flies *n* = 76/73; and ticks *n* = 39/44. When there was more than one ectoparasite specimen available per ectoparasite family per bat, one specimen for sequencing was selected based on morphological similarity, sequencing one individual of each morphotype of each family. No haplotypes from bugs and most tick specimens matched existing sequences deposited in GenBank or BOLD at the species level (divergence > 4%), with the exception of one of the tick lineages showing similarity to *Carios kelleyi* (96.95% in GenBank, 97.07% in BOLD). Similarly for nycteribiid bat flies, there were no species level matches for neither COI nor 18S markers. In the case of streblid bat flies, 14 sequences matched (∼99-100% in GenBank-BOLD databases) with five known species for COI (see Table 2); but there were no matches for 18S. These marker differences are likely attributable to database coverage, since where reference sequences were available for COI and 18S from the same specimen/study, BLAST results for our sequences were consistent both markers. For brevity and to provide comparability with larger numbers of reference sequences further reporting of diversity, divergence statistics and taxonomic identity will be focused on COI results.

**Table 2.**
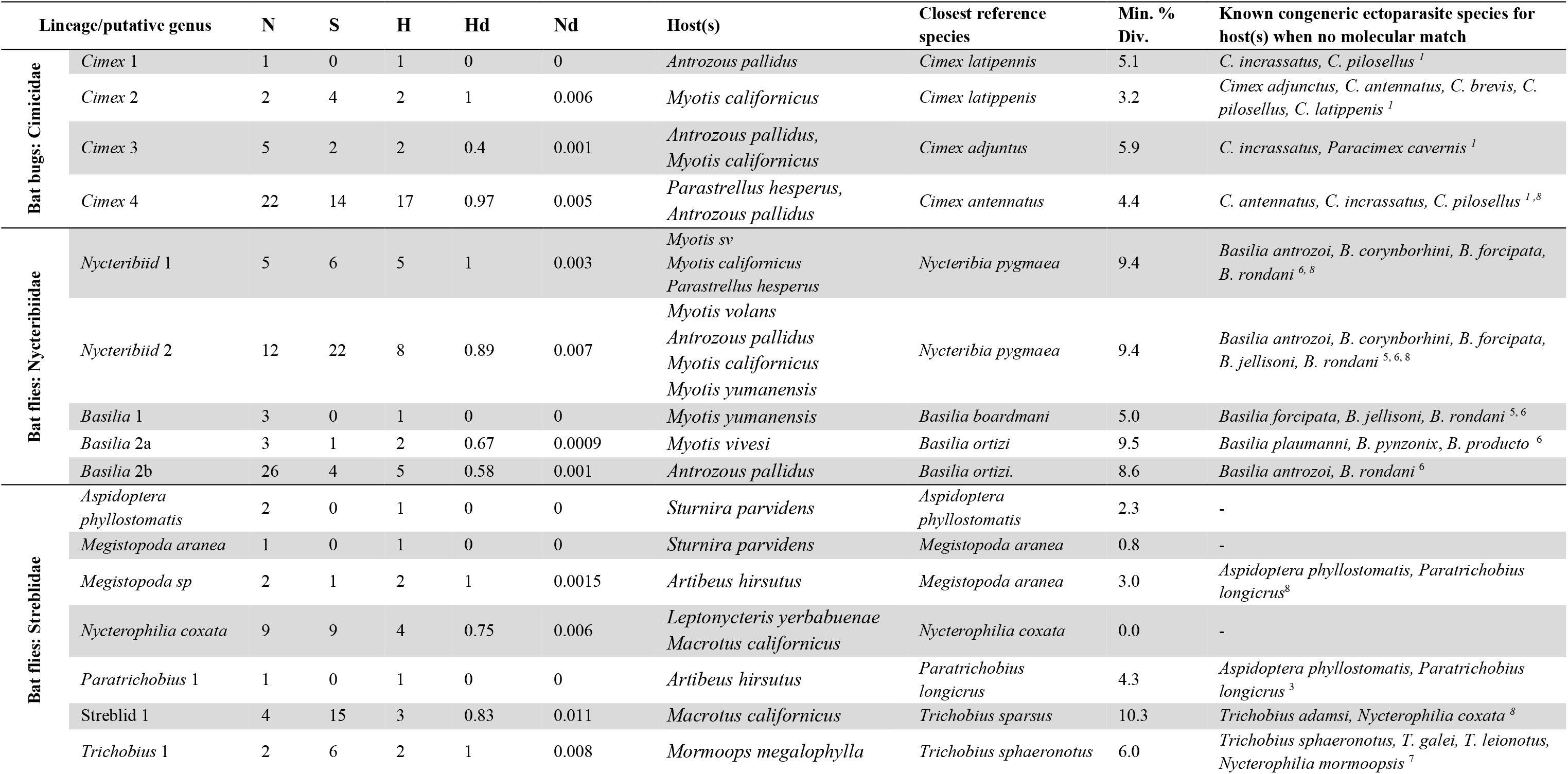
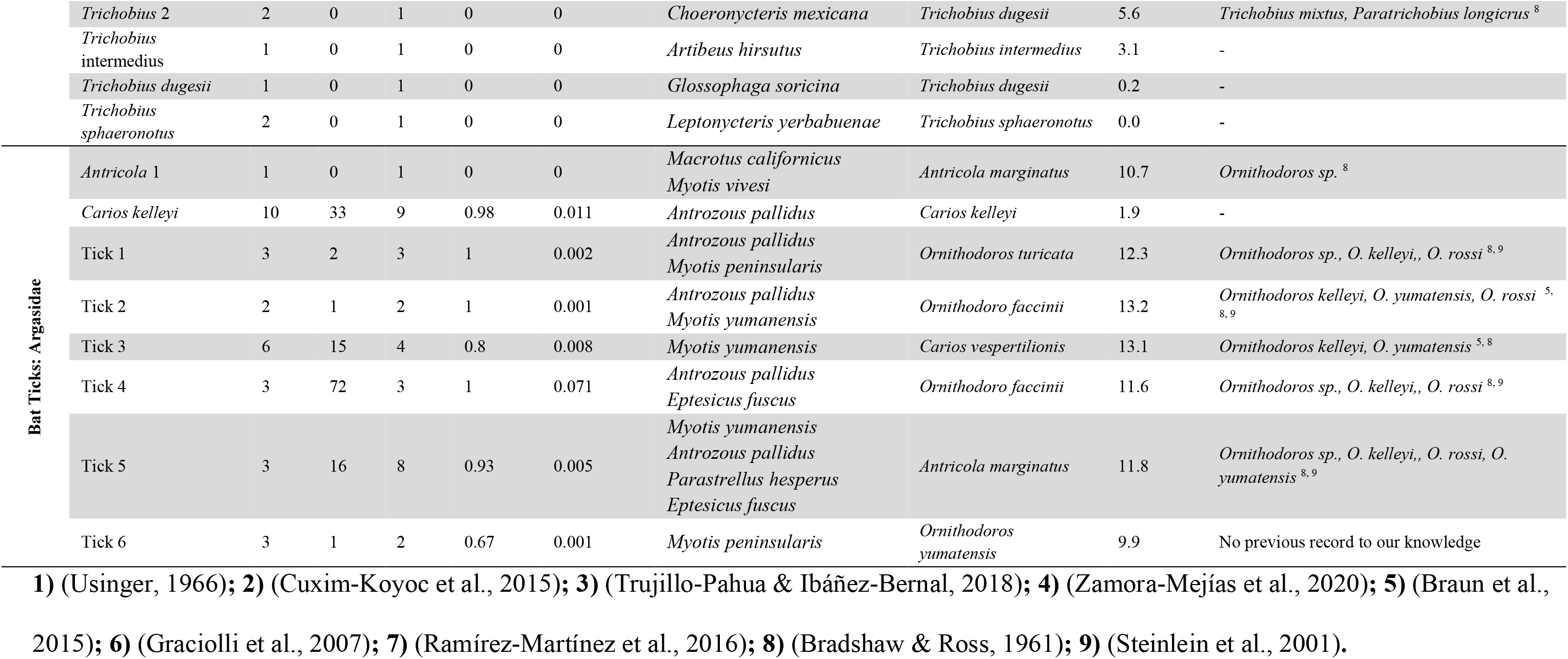
Molecular summary statistics for the ectoparasites lineages identified in this study. *N*, number of sequences within each lineage; *S*, number of segregating sites; *H*, number of haplotypes; *Hd*, haplotype diversity; *Nd*, nucleotide diversity, and Min. % Div., minimum percentage sequence divergence from closest reference sequence.

### Genetic diversity

Genetic diversity statistics for each COI lineage in each ectoparasite family are summarised in Table 2. Excluding four lineages represented by single individuals (e.g. *Cimex* 1), and three lineages including only individuals with the same haplotype (e.g. *Basilia* 1), nucleotide diversity (Nd) ranged from 0.001-0.006, with the highest value 0.071, presented by Tick 4 lineage, with three haplotypes (H) in three sequences (Table 2). In general, the number of haplotypes were close to or the same as the number of sequences tested per lineage.

### Phylogenetic assessment of Baja peninsula bat bug sequences

The best fit evolution model for the COI gene set was GTR+G+I, and K2+G+I for 18S. For each marker, 30 sequences were generated, forming four novel lineages with respect to reference sequences (Figure 3). Genetic divergence between the four peninsular lineages ranged from 9.9% to 17.1% (Appendix 4), and between 7.2% and 20.9% against reference sequences, with *C. latipennis, C. antennatus* and *C. adjuntus* presenting the lowest divergence values. Phylogenetically all the lineages sit within the *Pilosellus* complex (Talbot, Vonhof, Broders, Fenton, & Keyghobadi, 2016; Usinger, 1966), of North American members of the *Cimex* genus.

**Figure 3.**
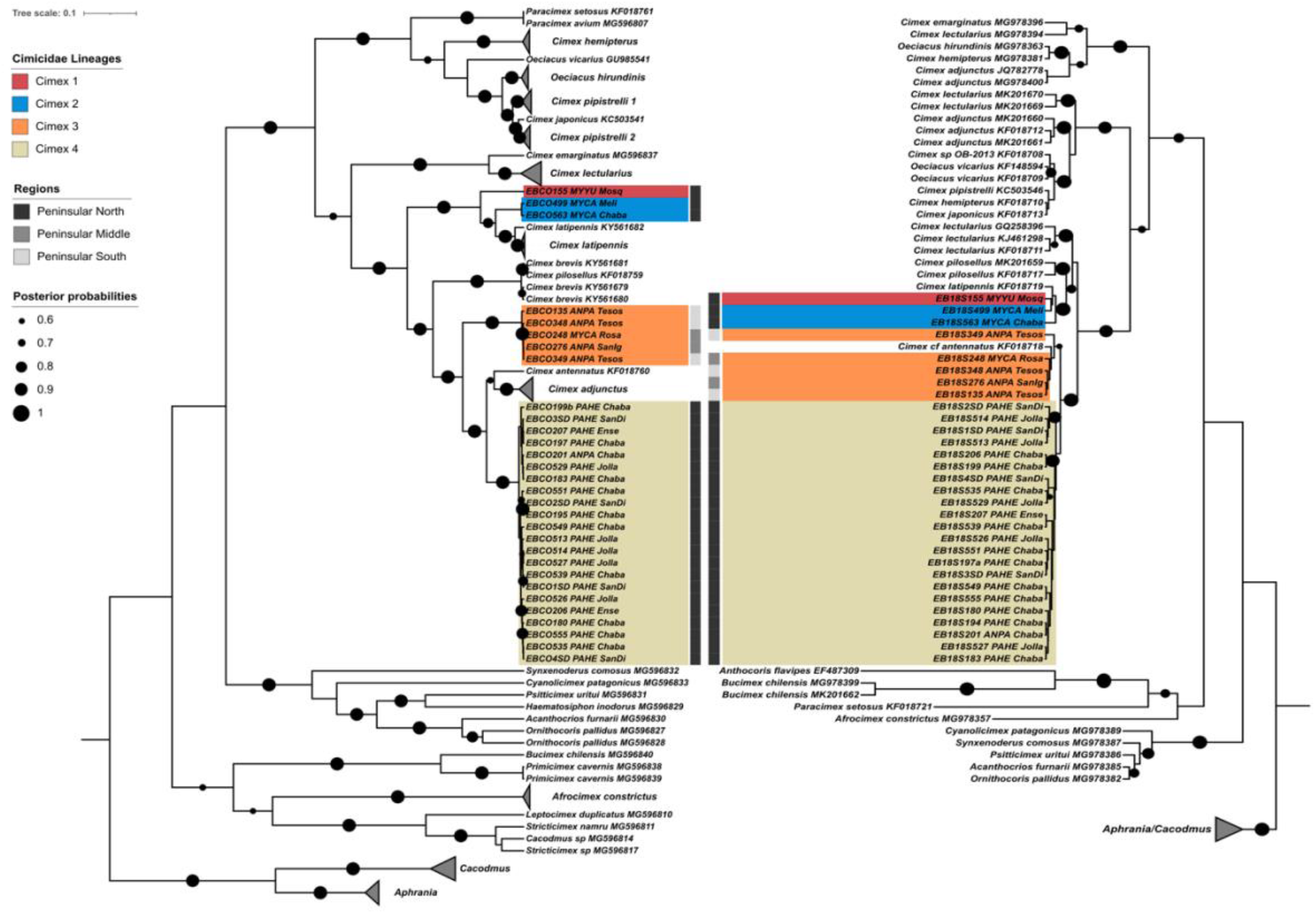
Bat bug phylogenetic trees obtained under Bayesian analysis using the mitochondrial COI (left) and the ribosomal 18S markers (right). Posterior probability support is indicated by the size of black circles at tree nodes. Where available, information for location and host is written next to each reference sequences label. To improve clarity of the tree, collapsed clades of reference sequences are shown as grey triangles.

For COI, *Cimex* 1 is represented by a single specimen (EBCO155) obtained from a *Myotis yumanensis* host at Mosqueda, and forms a sister lineage to *Cimex* 2, represented by two haplotypes from two bugs parasitizing *M. californicus* hosts, which were also sampled from northern sites. The 18S analysis also placed EB18S155 in a separate clade, but *Cimex* 2 is paraphyletic. Both markers placed *Cimex latipennis*, which is distributed from Canada to the north western USA (Usinger, 1966), as the closest named molecular reference species to these 2 lineages. *Cimex* 3 is represented by a monophyletic clade for COI, consisting of two haplotypes, sampled from specimens collected from four *Antrozous pallidus* and one *M. californicus* hosts, all distributed in the southern half of the peninsula (sites Rosarito, San Ignacio and Tesos). For 18S, a *C. cf antennatus* reference sequence nests within the clade with greater than 60% of posterior probability support. For COI, reference sequences for *C. pilosellus* and *C. brevis* formed a clade which appeared to be ancestral to *Cimex* 3, with 13% divergence (Table 3). There is no molecular reference for *C. brevis* in the 18S analysis, and *Cimex* 4 is the closest sister lineage.

The *Cimex* 4 COI lineage includes 17 haplotypes derived from 22 specimens, where 21 of the bugs were found parasitizing *Parastrellus hesperus* individuals, and one from *A. pallidus* (EBCO201/EB18S201), all from northern sites (Figure 1), with *C. antennatus* as the closet sequence match (9.3%; Figure 3), followed by *C. adjunctus*. Sequences of *C. adjunctus* in the 18S topology clustered in a clade with other *Cimex* species with mixed origins, including some non-Palearctic species. This may represent misidentification of those specimens, or mistakes in the annotation of sequences submitted to GenBank. *Cimex incrassatus* is reported as occurring on *A. pallidus* hosts (Usinger, 1966), but no reference sequences are available. Therefore, one of the novel bug lineages may represent this species, but further morphological assessments are required for confirmation.

### Phylogenetic assessment of bat fly sequences

The best fit sequence evolution models were GTR+G for the COI gene set, and T92+G+I for the 18S data. In total, 77 bat flies were sequenced, yielding 76 sequences for COI and 73 for 18S amplicons respectively. Individual sequences for two specimens for COI, and three for 18S did not pass sequence quality thresholds and were discarded. In the final sequence set for the COI marker, there were 49 sequences from the Nycteribiidae family (wingless bat flies) and 27 for the Streblidae family (winged bat flies), representing ten novel lineages and six new species records for Mexico. For the 18S marker, 46 sequences were generated for the Nycteribiidae family, and 26 sequences for the Streblidae family. For this marker, nine novel lineages were observed (18S sequences corresponding to the Basilia 2a COI lineage could not be amplified), along with six new species records for Baja.

In the COI phylogenetic analysis, the 49 Nycteribiidae sequences formed five lineages, all of which appeared to be novel with respect to GenBank references. Genetic divergence among Baja lineages ranged from 2.9 to 14.5%, and up to 16% against reference sequences (Appendix 5). For COI, Nycteribiid 1 and 2, formed sister clades with 4.4% divergence, and 10% divergence from the closest reference haplotypes derived from Asian *Nycteribia*, species. For 18S the closest references to Nycteribiid 1 and 2 were North American species not available for COI, *Basilia corynorhini* and *Basilia forcipata*, respectively. Nycteribiid 1 and 2 lineages were primarily associated with *Myotis* bat hosts, but with one Nycteribiid 2 haplotype recovered from a *Parastrellus hesperus* host in the 18S dataset (specimen EN18S514, Figure 4).

**Figure 4.**
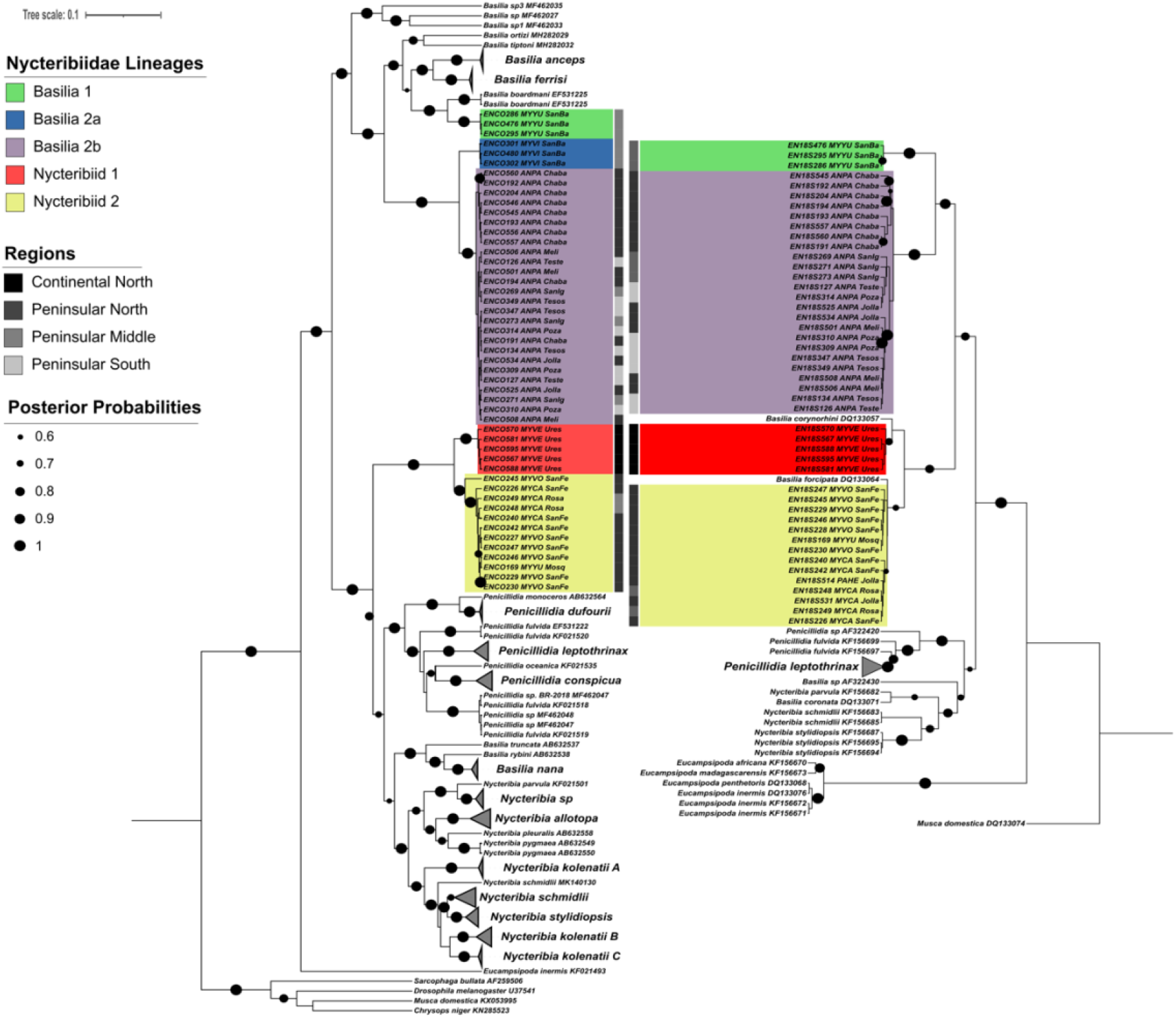
Bat flies of the family Nycteribiidae phylogenetic tree obtained under Bayesian analysis using the mitochondrial COI (left) and the ribosomal 18S markers (right). Posterior probability support is indicated by the size of black circles at tree nodes. Where available, information on location and host is written next to each reference sequences label. For clearer visualization of the tree large clades of reference sequences collapsed grey triangles.

The three other Baja lineages formed clades associated with *Basilia* reference sequences from species recorded in Madagascar, USA and Panama. *Basilia* 1 has 4.9% divergence from *Basilia boardmani*, a bat fly distributed throughout the United States parasitizing *Myotis* bats (Graciolli et al., 2007). Specimens with *Basilia* 1 haplotypes were sampled at mid-peninsula, parasitizing *Myotis yumanensis*, which is widely distributed throughout western North America (Braun, Yang, González-Pérez, & Mares, 2015).

Genetic divergence among lineages *Basilia* 2a and 2b was 2.9% (Table 2), representing the threshold for intra/inter interspecific values (Hebert et al., 2003; Ratnasingham & Hebert, 2013). We assign them as distinct lineages due to their different host species, where *Basilia* 2a parasitized *M. vivesi*, and *Basilia* 2b appeared to be restricted to *Antrozous pallidus* (Figure 4). The spatial distribution of *Basilia* 2b haplotypes across the peninsula, suggests potential long distance co-dispersal with their hosts *A. pallidus* (Figure 4), providing evidence supporting bat movement along Baja. *Basilia* 2b may represent *B. antrozoi*, which has been previously reported to parasitize *A. pallidus* (Graciolli et al., 2007).

The 11 Streblidae lineages had genetic divergence for COI of 0.01% to 18.9% for sequences sampled within this study, and up to 21.6% against their reference sequences (Appendix 6). For the five novel lineages from this family four were named according to the closest genus in the phylogenetic analysis (*Trichobius* 1, *Trichobius* 2, *Paratrichobius* 1 and *Megistoposa sp.,* Figure 5), with genetic divergence values ranging from 3% to 8.4% from each of the closest corresponding clades. The Streblid 1 lineage presented 10.8% divergence against its closest reference, *T. sphaeronotus* (Appendix 6, Figure 5), and is therefore not attributed to an existing genus.

**Figure 5.**
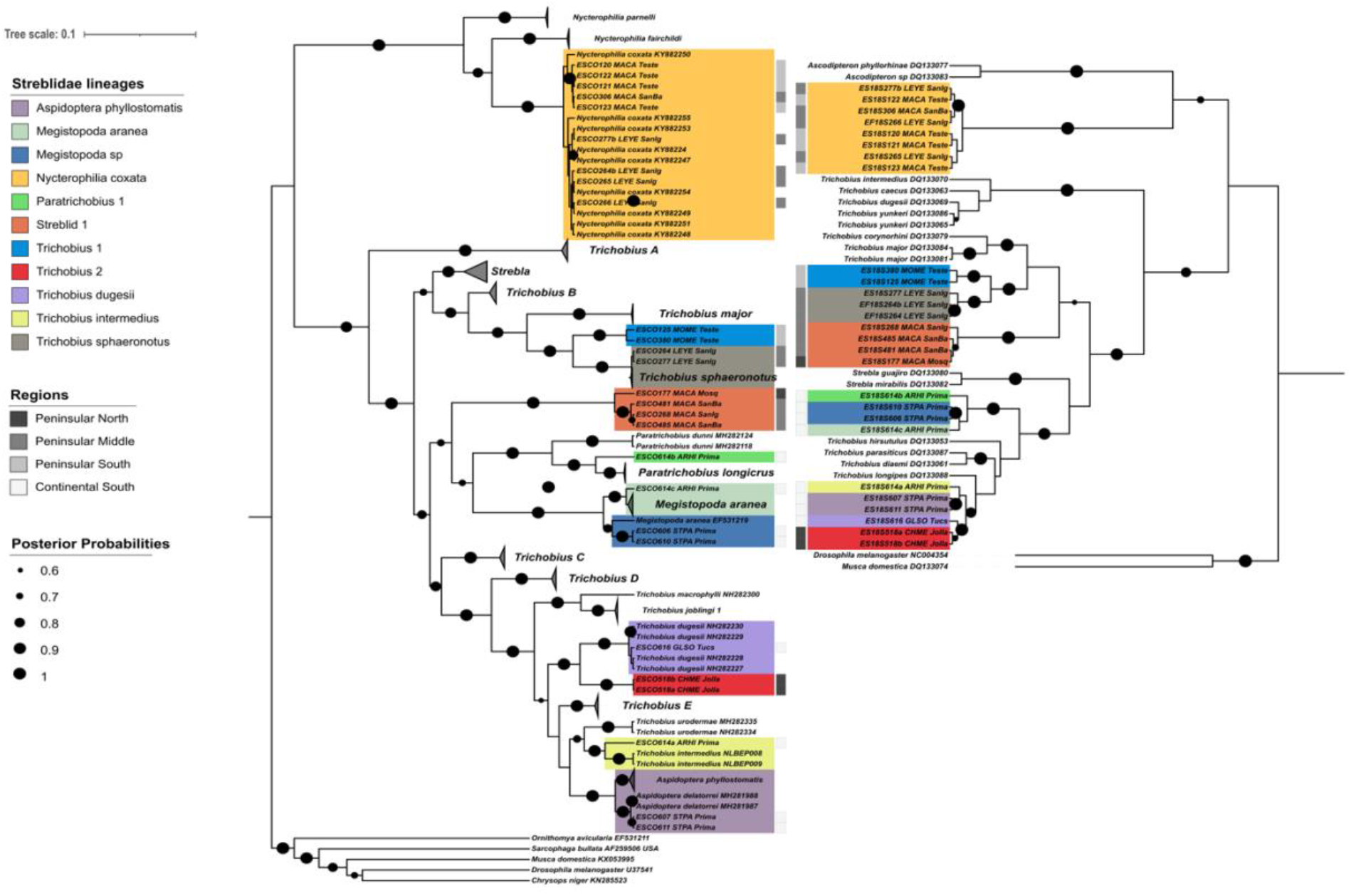
Bat flies of the family Streblidae phylogenetic tree obtained under Bayesian analysis using the mitochondrial COI (left) and the ribosomal 18S markers (right). Posterior probability support is indicated by the size of black circles at tree nodes. Where available, information on location and host is written next to each reference sequences label. For clearer visualization of the tree large clades of reference sequences collapsed grey triangles.

The other streblid clades matched sequences from six known species: *Aspidoptera phyllostomatis, Megistopoda aranea, Nycterophilia coxata, Trichobius dugesii, T. intermedius* and *T. sphaeronotus* (Figure 5), representing new records for these species in Baja and western Mexico. For the specimens classified as *N. coxata, T. dugesii, T. intermedius* and *T. sphaeronotus* (Figure 5, Appendix 6), there was less than 1.5% against the GenBank references. The fly lineages *A. phyllostomatis* matched with two reference sequences classified as *A. delatorrei* (Figure 5). However, *A. phyllostomatis* and *A. delatorrei* reference sequences only differed by 2.1 - 2.3%, which falls at the threshold for species level differentiation. Without further information on the source *A. delatorrei* specimens, we decided to retain the *A. phyllostomatis* classification for our specimens. In the case of the *M. aranea and M. sp* lineages, there were two separate clades of *M. aranea* reference sequences present. The *M. sp* specimens (ESCO606 and ESCO610, dark blue clade, Figure 5) were grouped closer to *M. aranea* EF531219 reference sequence, with 2.9% and 3.1% divergence, suggesting a different species from the *M. aranea* EF531219 reference, as well as from the other *M. aranea* clade (bright blue clade, Figure 5), with 4.4% genetic divergence. Supporting this, it is noted that each lineage had different host species, *Sturnira parvidens* and *Artibeus hirsutus*, respectively (Table 2). Most streblid lineages were parasitizing fruit-nectar feeding bats (*Phyllostomidae*) over the mid and northern peninsula, with the exception of *Trichobius* 1 found on *Mormoops megalophylla* hosts, (family *Mormoopidae*), which feeds on insects. *Trichobius* 1 was the only streblid fly lineage found in the south of Baja. The other fly lineages were distributed in the southern continental sites (Tucson and Primavera sites, Figure 1, Table 1). Outside of the closest sequence matches, none of the other species reported as parasitizing hosts in Baja (Table 2) appear to have reference sequences in GenBank. Genetic divergence within groups from the lineages in this study showed levels between 0.0% and 1.1%, with the exception of *A. phyllostomatis*, which showed a 3.3% divergence within its group (Appendix 6).

### Phylogenetic assessment of Baja peninsula bat tick sequences

The best fit sequence evolution models were GTR+G+I for the COI gene set, and K2+G+I for the 18S gene set. There were 45 ticks sequenced from the Argasidae family (soft ticks), with 39 sequences generated for COI, and 44 for 18S, where six specimens failed to amplify for COI (specimens ETCO157, ETCO306, ETCO338, ETCO480b, ETCO480c and ETCO485), and one specimen failed to amplify for 18S (specimen ET18S457). One new Baja species record and seven potential novel lineages were obtained (Figure 6). The only match with GenBank and BOLD COI records (95.95% and 97.07%, respectively) was the soft tick *Carios kelleyi*, for 11 specimens (parasitizing *A. pallidus* hosts) with intra-clade divergence of 1.94% to 2.47% (Figure 6, Appendix 7). For COI data, *Antricola* 1 lineage had 10.6% divergence from *Antricola marginatus*, and 14.6% from *A. mexicanus* (Appendix 7). *Ornithodoros faccini* had a divergence of around 11% for clades *Carios kelleyi*, Tick 1 to 5, and *O. yumatensis* showed divergence of around 9.9% with respect to Tick 6 lineage. Ticks of lineage Tick 6 were recovered from *M. peninsularis* from which there are no previous records of bat ticks to our knowledge.

**Figure 6.**
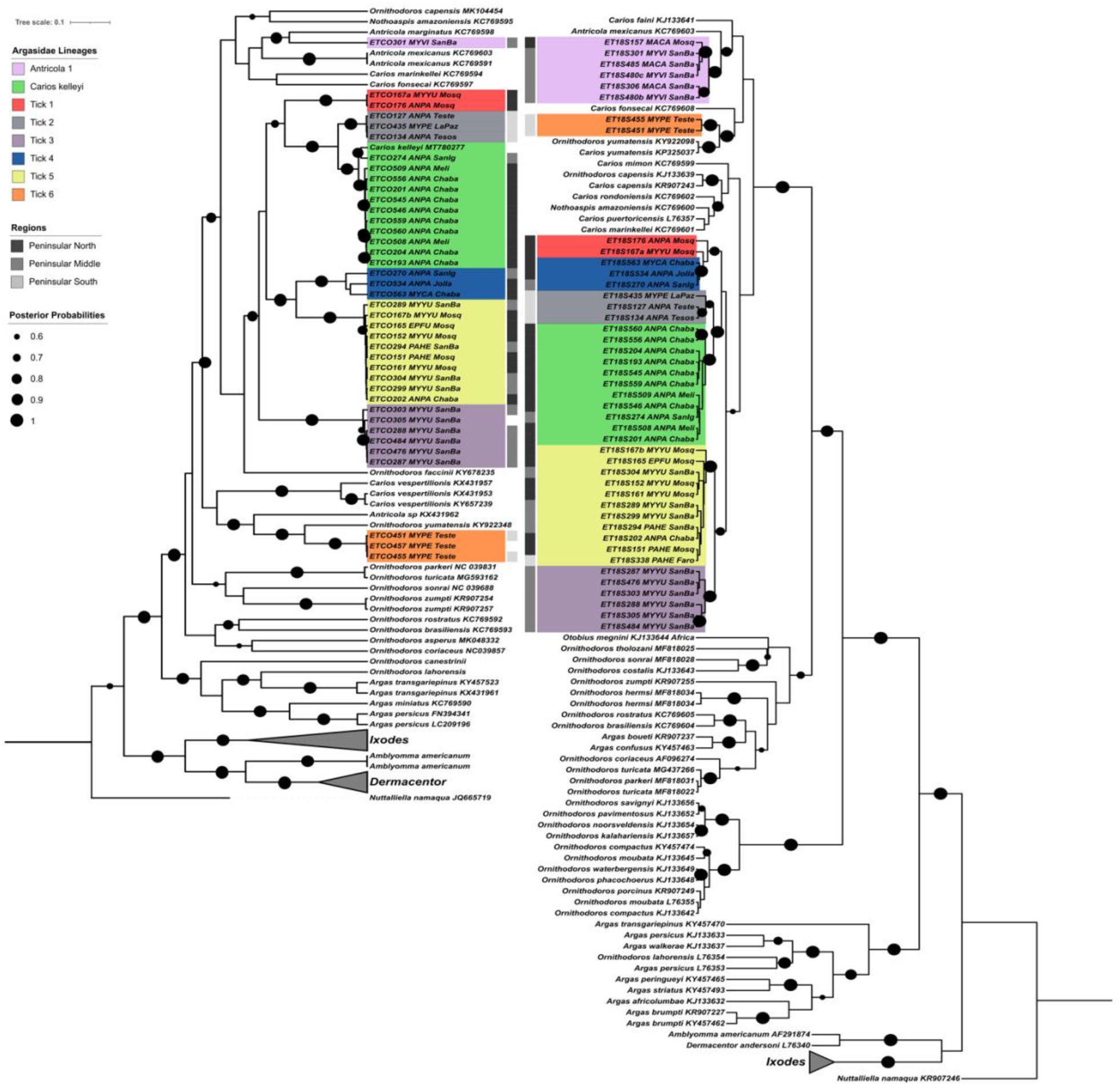
Bat ticks of the family Argasidae phylogenetic tree obtained under Bayesian analysis using the mitochondrial COI (left) and the ribosomal 18S markers (right). Posterior probability support is indicated by the size of black circles at tree nodes. Where available, information on location and host is written next to each reference sequences label. For clearer visualization of the tree large clades of reference sequences collapsed grey triangles.

All sequences from this study generated the same lineages for both COI and 18S genes (Figure 6). Lineages *C. kelleyi* and Tick 1 to 5 formed a monophyletic group with respect to the reference sequences with posterior support greater than 0.6 for both markers, while *Antricola* 1 and Tick 6 lineages were positioned in separate clades. The topology of sister clade relationships varied slightly between markers, particularly around deeper nodes which had posterior probability support less than 0.85. This suggests more data is required to resolve deeper taxonomic relationships among species. *Antricola* 1 and Tick 6 were separated around shallow deep nodes in both analyses, where the absence or presence of reference sequences influenced their topological proximity (Figure 6). None of the reference sequences from *Antricola*, *Carios*, and *Ornithodoros* genera, form monophyletic groupings with respect to genus nomenclature.

Tick lineages 2 and 6 (grey and orange, Figure 6) were only observed in the south of the peninsula, with hosts *A. pallidus* and *M. peninsularis*, and *M. peninsularis* respectively. These two species of bats share at least one confirmed roosting site within the study region, at Tesos (Figure 1), suggesting a potential interchange of host species for Tick lineage 2. Evidence of singular host specificity was observed for the lineages *C. kelleyi* (*A. pallidus*), Tick 3 (*M. yumanensis*), and Tick 6 (*M*. *peninsularis*), all observed exclusively on the same hosts across sites and field seasons. The *Antricola* 1 lineage (Figure 6, lilac clade, ETCO301_MYVI,) was recovered from one specimen using the COI marker, and grouped with *Antricola marginatus* (found in the South East of Mexico), followed by *A. mexicanus*. In the 18S topology, five additional sequences were obtained, grouped with *A. mexicanus*, as no *A. marginatus* reference was available. Two host species for ticks of this lineage were identified, *M. vivesi* and *Macrotus californicus*, sampled mostly at mid-peninsula (Figure 1 and Figure 2). Tick 5 lineage had the most diverse host and spatial distribution, being found on four bat species, and at sites from the north to the south of the peninsula (yellow clade, Figure 6).

## Discussion

This study represents the first molecular characterisation and phylogenetic analysis of bat bug, bat fly and bat tick diversity along the Baja California peninsula and northwestern Mexico, a region where the bat ectoparasite fauna is largely undescribed. From a total of 292 ectoparasites specimens, from 17 bats species, evaluated for COI and 18S markers, 21 novel genetic lineages, plus seven new species records for the region, were found. Some of the novel lineages may derive from previously recorded species with no reference sequences available, while others are likely to represent new species. Overall the work demonstrates that the northwestern region of Mexico hosts a high diversity of previously unknown bat ectoparasites.

### Bat bugs

Four novel lineages of *Cimex* bugs were identified. The lineages showed evidence of host preference, being found primarily on *M. yumanensis*, *M. californicus*, *A. pallidus* and *P. hesperus*, respectively for lineages 1-4. A threshold of 4% sequence divergence is typically taken as representing species level differentiation for COI in arthropods (Hebert et al., 2003), indicating Cimex 1, 3, and 4, could be classed as novel species under this criterion. Genetic divergence of *Cimex* 2 compared with *C. latippenis* was 3.2%, but *C. latippenis* has not previously been recorded parasitizing *M. yumanensis* (Braun et al., 2015), which might also suggest *Cimex* 2 as a potential new cryptic species.

Cimicid bugs have low inherent dispersal capacity, generally feeding for a few days, before dropping from the bat host to digest the blood in the roost, where they can survive without feeding for approximately 1.5 years (Ossa et al., 2019). The population structure of bat bugs is mainly influenced by bat movements (Ossa et al., 2019; Talbot et al., 2016; Talbot, Vonhof, Broders, Fenton, & Keyghobadi, 2017; Usinger, 1966). While the current data indicate the new *Cimex* lineages may have distinct regional distributions within Baja, their host species ranges extend further through western North America, suggesting these bugs may also have wider distributions, or come into contact with hosts which migrate over long distances at roost sites. For example, *M. yumanensis* individuals captured in the northern peninsula in our study had mitochondrial haplotypes matching reference sequences from bats sampled in Alaska (data not shown). This raises the possibility of long-range mixing of microbial pathogen communities in bat bugs along the west coast of North America.

### Bat flies

Ten novel genetic lineages and six new records of bat flies for the study region were found. Nycteribiid flies were more abundant in the northern temperate sites, while streblids were more abundant in the southern and subtropical sites, supporting trends noted by Dittmar *et al*. (Dittmar et al., 2006). To our knowledge, *Nycteribia* species have not previously been reported for bats with ranges in Baja. Streblid flies were present on bats from the Phyllostomidae, which in general are fruit and nectar feeders, with the exception of the insectivorous *Macrotus californicus*; as well as *Mormoops megallophyla*, from the family Mormoopidae. Nycteribiids parasitized only bats Vespertilionid bats, which include insectivores and omnivores.

In previous studies, Nycteribiid bat flies have typically been found to be host specific (de Vasconcelos, Falcão, Graciolli, & Borges, 2016; Dittmar et al., 2006; Saldaña-Vázquez et al., 2019), with some exceptions (Olival et al., 2013; Wilkinson et al., 2016). We found a similar pattern, with most Nycteribiid lineages exhibiting host specificity, despite having hosts with overlapping ranges and which may share roost sites (e.g. *Myotis* species in northern Baja). The Nycteribiid 2 lineage was found on multiple *Myotis* species, which could potentially indicate specificity at a genus level. Streblid winged flies have been described as mostly non host-specific (Dittmar et al., 2006). In our data *N. coxata* was found parasitizing *Macrotus californicus* and *Leptonycteris yerbabuenae*, both Phyllostomidae species, which are known to share roost site in Baja (Álvarez-Castañeda et al., 2015), implying potential horizontal transmission.

Lineages *Basilia* 2a and *Basilia* 2b (COI divergence 2.9%) are restricted to hosts *Myotis vivesi* and *Antrozous pallidus* respectively. *M. vivesi* is endemic to the Gulf of Cortes, and restricted to coastal habitats because of its piscivorous diet (Blood & Clark, 1998; Herrera-Montalvo, Flores-Martínez, & Sánchez-Cordero, 2017). *A. pallidus* is sympatric with *M. vivesi* on the Gulf coast but has a wider distribution across western North America. It is primarily an insectivorous feeder, but also includes scorpions and nectar in its diet (Frick, Hayes, & Heady, 2009). *Basilia* fly records previously reported for *M. vivesi*, but without reference sequences, include *B. plaumanni, B. pynzonix,* and *B. producto* (Graciolli et al., 2007), while flies parasitizing *A. pallidus* have been described as *B*. *antrozoi* (Table 2). The threshold level of divergence between the *Basilia* 2a and *Basilia* 2b lineages may indicate recent divergence from a common ancestor, and potential incipient speciation driven by association with sympatric but ecologically differentiated hosts.

### Bat ticks

A new record of *Carios kelleyi* and seven novel tick lineages belonging to the Argasidae family were found. For both COI and 18S tick lineages 1-5 and *Carios kelleyi* formed a clade with more than 0.6 posterior support, suggesting they form a Baja or western North America endemic lineage of bat ticks derived from a common ancestor.

The genera *Antricola, Carios*, *Nothoaspis* and *Ornithodoros* are associated with bats and their roosting sites in Mexico (Sánchez-Montes et al., 2016). Compared to Ixodidae, the family Argasidae has few molecular studies and reference sequences (Porter & Hajibabaei, 2018). Classifications at genus level are controversial, with many genera being paraphyletic in existing phylogenies, and debates over synonymous use of genus names such as *Carios* and *Ornithodros* (Burger et al., 2014). Such ambiguities are also reflected in our phylogenies. Further, while many key internal nodes had posterior support greater than 0.6, differences in reference sequence availability made it challenging to interpret the consistency of inter-clade placement between makers.

Argasid ticks have previously been reported for the bat species in this study, primarily *Ornithodoros* species (Table 2), but with the exception of *Carios* (*Ornithodros*) *kelleyi*, none are close molecular matches for our sequences. In ticks, for COI, the threshold for between genus divergence is considered to be above 10% (Hebert et al., 2003). The observed COI divergence among lineages of this study (6.1% up to 19.3%), and to reference sequences (9.9% to 22.8%), suggests that the Baja lineages could represent novel species and potentially novel genera.

We found apparent host-specific and generalist lineages for the ticks reported here. Lineages that appeared host-specific were restricted to single sites, while generalists were found across multiple sampling locations. For example, Tick 3 parasitizing *M. yumanensis* was found only in San Basilio, and Tick 6 was found only on *M. peninsularis* sampled in Teste. *Carios kelleyi* was found to only parasitise *A. pallidus* in this study, and was found at three sites in the mid and north peninsula. However, *C. kelleyi* is known to parasitise multiple species across North America and Cuba (Gill et al., 2004), and the reference COI sequence used here derives from an *Eptesicus fuscus* bat sampled in New Jersey, eastern USA (GenBank accession code: MT780277). Argasid ticks show a continuum of hosts-specificity (Cumming, 1998; Esser et al., 2016), but tick stage-cycle must be considered, with immature ticks being more generalist than adult conspecifics (Esser et al., 2016; Nava & Guglielmone, 2013). In this study, life stage was not assessed while collecting ticks, therefore it is possible that there are gaps in host range regarding unidentified larvae that were not sequenced.

The Tick 5 lineage was found on multiple species in the mid and north peninsula, but the only specimen recorded for the south peninsula was collected from *Parastrellus hesperus* (specimen 338, Faro site, 18S only). Since *P. hesperus* is rare in southern *Baja*, the presence of this tick indicates potential dispersal of *P. hesperus* from the north to the south peninsula.

### Limits on ectoparasite sampling and identification

The present study identified 21 novel genetic lineages, plus 7 new ectoparasite species records, from 138 bats of 17 species, sampled across 18 sites and 2 years. This suggests a diverse ectoparasite fauna in this previously unsurveyed region of Mexico, but it is also likely to be an underestimate of the true diversity, due to constraints around sampling effort. Bat sampling was limited to May-August and conducted at relatively accessible locations with water sources to facilitate bat capture. While our sampling sites were chosen to be representative of habitat types across Baja, increasing the spatial and temporal scope of sampling would be likely to increase the number of ectoparasite species discovered. Assessment of ectoparasite fauna found in roosting sites against those found feeding directly from their hosts and expanding seasonal coverage will be important for future work.

Although previous studies report limited data on bat ectoparasites from North Western Mexico and South-Western USA (Bradshaw & Ross, 1961; Braun et al., 2015; Pérez et al., 2014; Usinger, 1966), they do not integrate morphological and molecular information. For many species, no reference sequence is available from voucher specimens, or there are errors in species identifications and incorrect annotation of reference sequences. Therefore, for all the ectoparasite groups in this study, further work is needed to unify molecular and morphological characterisation, to fully confirm which lineages represent previously undescribed species, and which are species with morphological descriptions but no previous molecular record. This is particularly important for bat tick lineages where input from expert morphologists is needed to account for different life stages.

## Conclusions

This work presents an initial description of bat ectoparasite diversity relevant to western North America, providing resources useful for future ectoparasite surveys and studies of host-ectoparasite ecology, and evaluation of zoonotic disease risks. Future work should focus on expanding spatial and bat host species coverage, integrating morphological and molecular characterisation of ectoparasite species, profiling ectoparasite microbiomes and viromes, and understanding the ecological and environmental factors that influence host-parasite community structure and evolution. Bat populations and habitat in Baja California are vulnerable to anthropogenic pressures, and such knowledge will be vital for informing assessments of population status and extinction risks of both hosts and parasites.

## List of abbreviations

18S: 18S ribosomal DNA
BCP: Baja California peninsula
BY: Bayesian analysis
COI: Cytochrome Oxidase subunit I.

## Declarations

### Ethics approval

All bat handling was carried out under the approval of the ethics committee of the Faculty of Biological Sciences, University of Leeds (AWCNRW170615); and following the Guidelines of the American Society of Mammalogists (Sikes *et al*., 2016). Sampling was carried out under the permits SEMARNAT-DGVS-008972-16 and SEMARNAT-DGVS-001642-18 issued by Secretaría del Medio Ambiente y Recursos Naturales (SEMARNAT) in Mexico. The latter included two *Myotis* bat species listed in the Mexican Official Norm for the protection of native species of flora and fauna in Mexico (NOM-059-SEMARNAT-2010, (Secretaría de Medio Ambiente y Recursos Naturales, 2010)), under the Pr (under special protection) and P (in danger of extinction) categories (*M. evotis* and *M. vivesi*, respectively); and sampling on protected reserves. When required, permission was also solicited and granted from private land owners. All samples were imported into the UK under permission of the Department for Environment, Food and Rural Affairs (DEFRA) from the Animal and Plant Health Agency (permit ITIMP19.0036). Ethical approval for bat capture, handling and sampling was granted by The Animal Welfare and Ethical Review Committee (AWCNRW 170615) from the University of Leeds.

### Availability of data and material

Sequences for the haplotypes reported in this study are available under GenBank accession numbers XXXX. (Data will be made available upon acceptance of the manuscript)

### Authors’ contributions

LANC and SG conceived study; LANC, AK, TK and SJG collected the material; DS donated samples; LANC performed DNA extractions and laboratory analyses; LANC analysed the data and wrote the first draft of the manuscript; SJG and LANC edited the draft to produce the final submission. All authors read and approved the submitted manuscript.

## Acknowledgements

We thank Dr. Sergio Ticul Alvarez Castañeda for his help with field logistics and for sharing his vast knowledge of bat distribution and behaviour over the peninsula; and to Dr. Alfredo Ortega and technician Izmene Rojas, all from The Northwest Centre of Biological Research (CIB), for their support and collaboration throughout this project. We also thank Dr. Winifred Frick for recommending site locations for sampling bats plus other vital information and contacts for this project; Alan Harper, Paul Heady and Quintin Frick for their invaluable help with fieldwork; to Dr. Eduardo F. Aguilera Miller for his help over the 20016-17 field seasons as well as Alejandro Najera, Halley O’Connor and Natasha Kebala for the 2018 season. Finally, thanks to Kieran Neal for support with some DNA extractions and generation of some PCR amplicons; and to Dr Michael Siva-Jothy for providing bed bug samples used for optimisation of PCR conditions and positive controls. We thank CONACYT and the Rufford Foundation for the funding provided, as well as the University of Leeds for all the resources provided for this research.

## Appendices

**Appendix 1.**
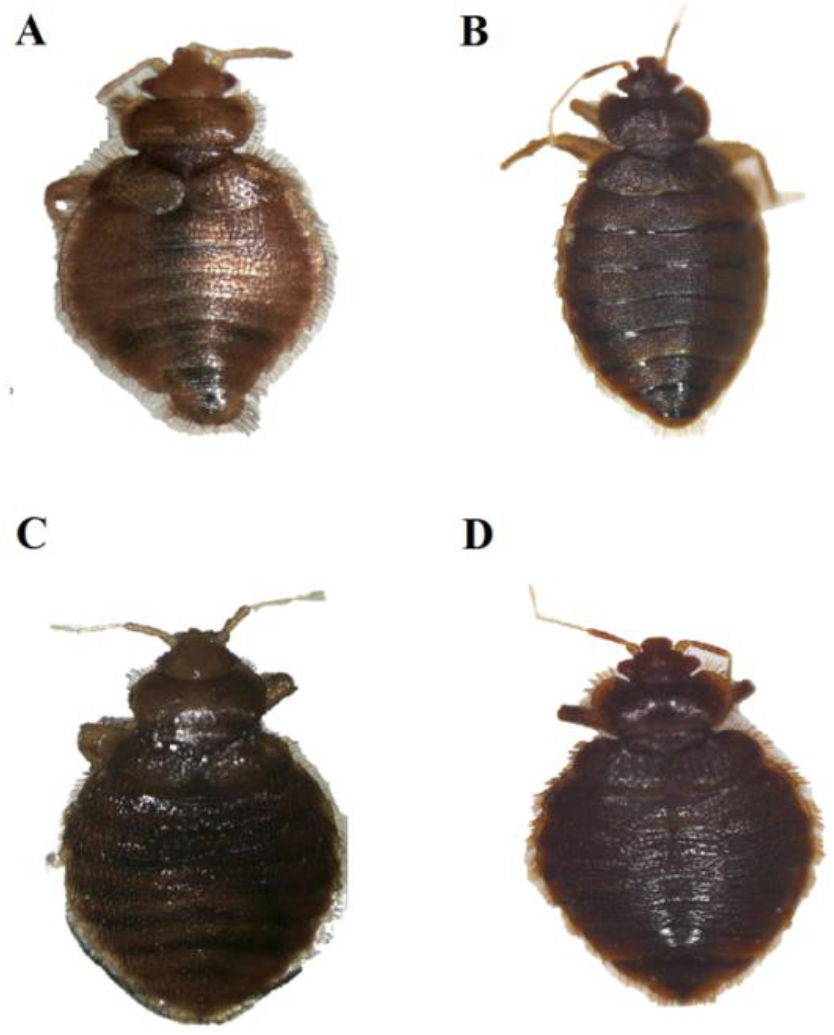
Pictures of a representative specimen (independent of sex determination) of each bat bug lineage: A, *Cimex* 1; B, *Cimex* 2; C, *Cimex* 3; and D, *Cimex* 4.

**Appendix 2.**
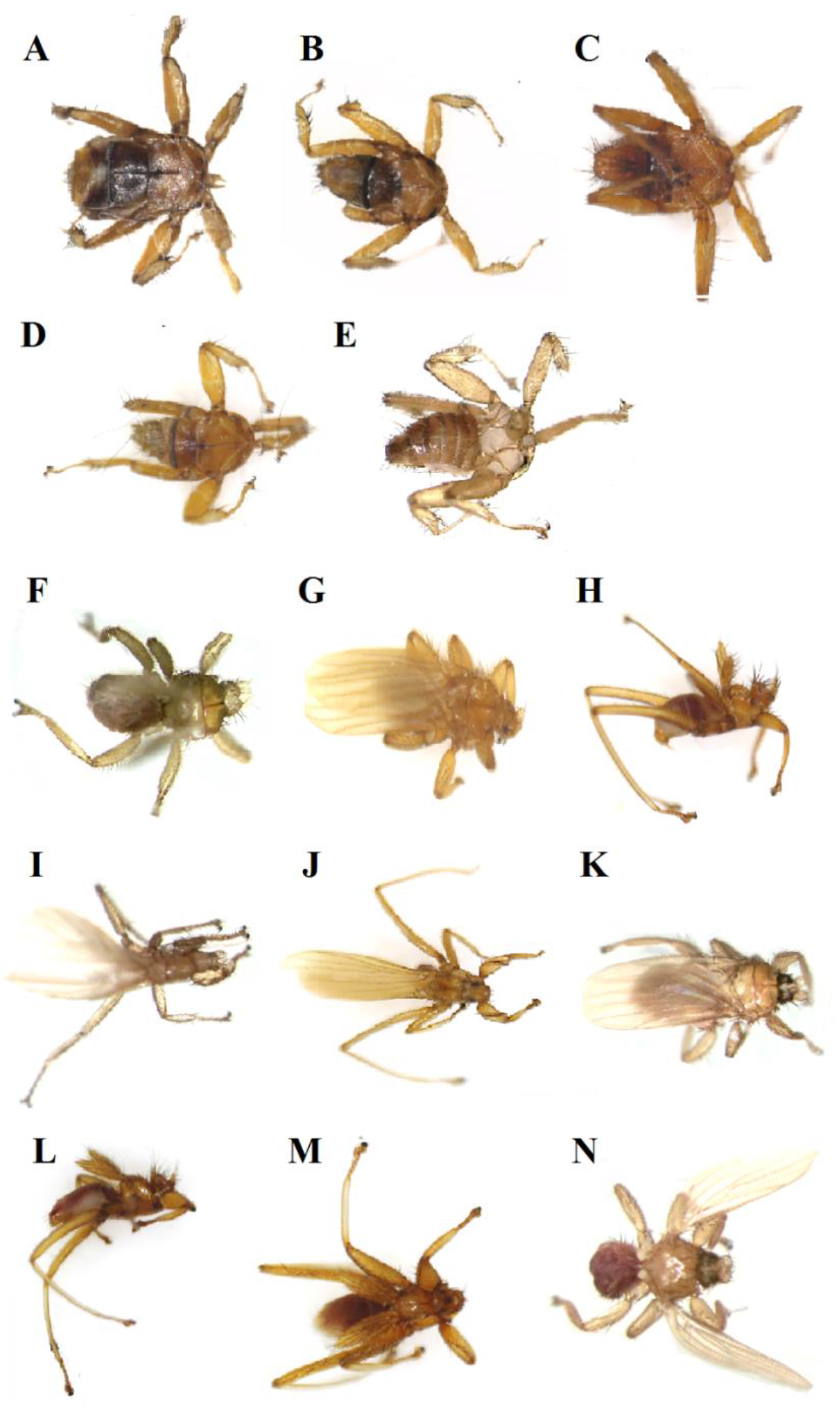
Pictures of a representative specimen (independent of sex determination) of each bat fly lineage: A, *Basilia* 1; B, *Basilia* 2a; C, *Basilia* 2b; D, Nycteribiid 1; E, Nycteribiid 2; F, *Aspidoptera phyllostomatis*; G, *Megistopoda aranea*; H, *Nycterophilia coxata*; I, *Paratrichobius* 1; J, *Trichobius* 1; K, *Trichobius* 2; L, *Trichobius* 3; M*, Trichobius sphaeronotus* 1; and N, *Trichobius dugesii*.

**Appendix 3.**
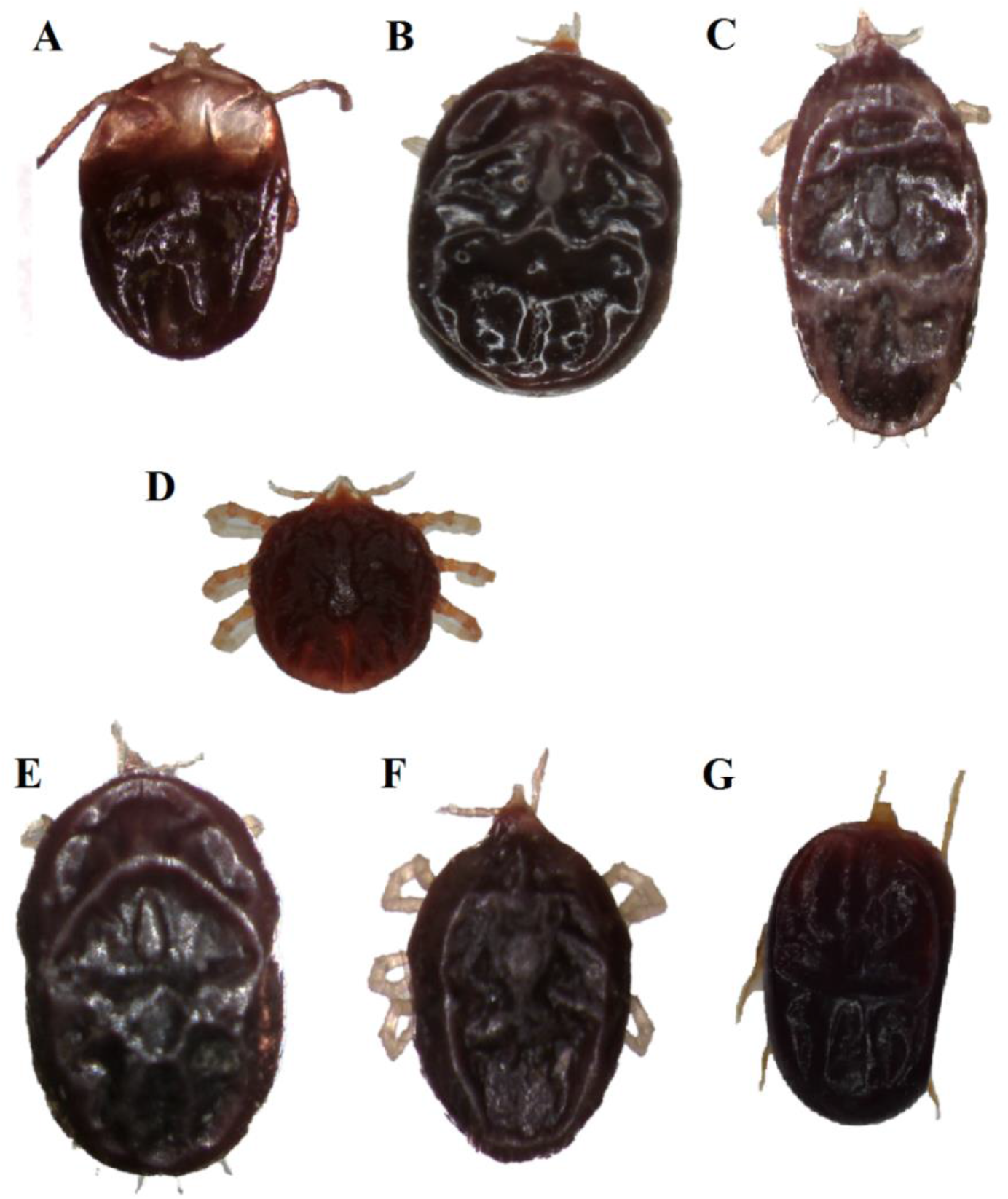
Pictures of a representative specimen (independent of sex determination) of each bat tick lineage: A, *Antricola* 1; B, *Carios kelleyi;* C, Tick 1; D, Tick 3; E, Tick 4; F, Tick 5 and G, Tick 6. There was not specimen available of Tick 2 lineage after sequencing.

**Appendix 4.**
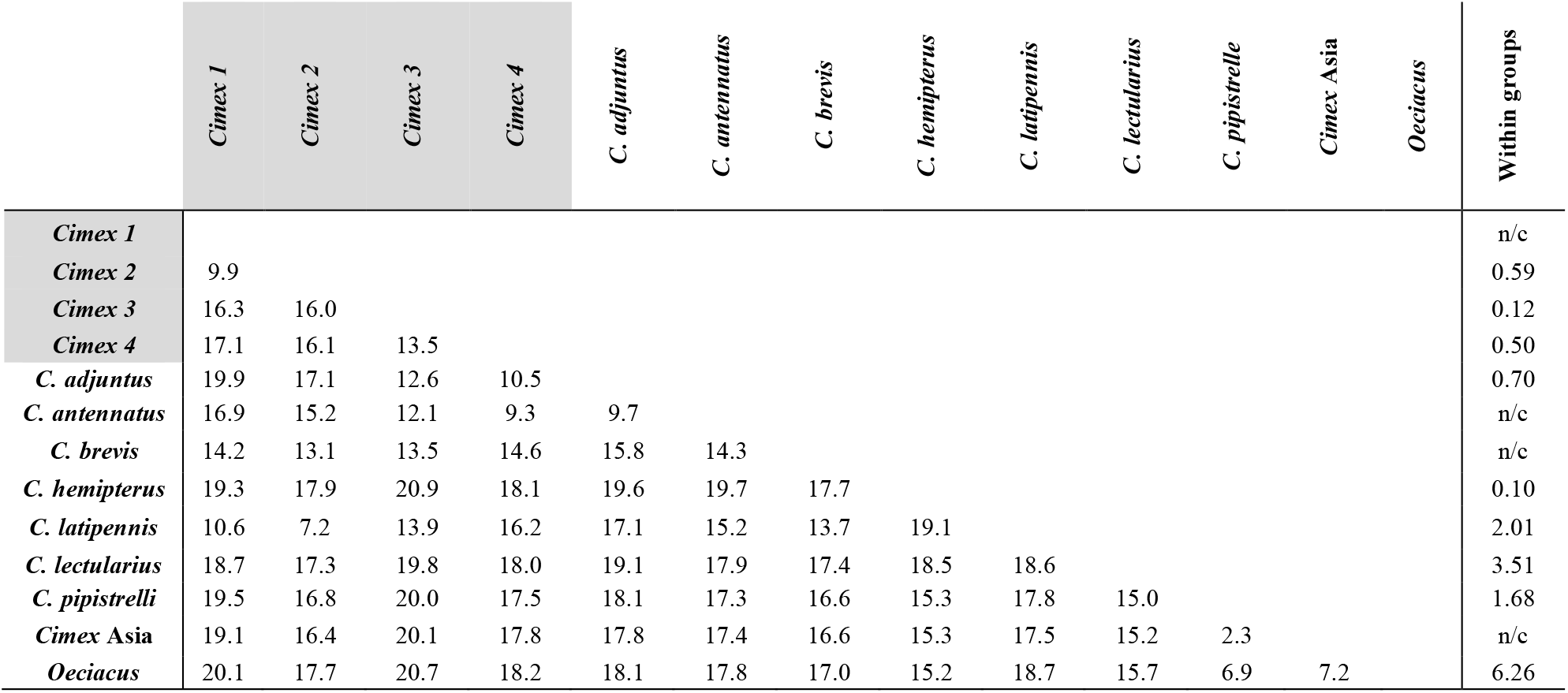
Estimates of percentage divergence (COI) over sequence-pairs between groups (diagonal left matrix), and within groups (most left column) of bat bugs (family Cimicidae), showing the lineages obtained in this study (shaded) versus closely related reference sequences from GenBank.

**Appendix 5.**
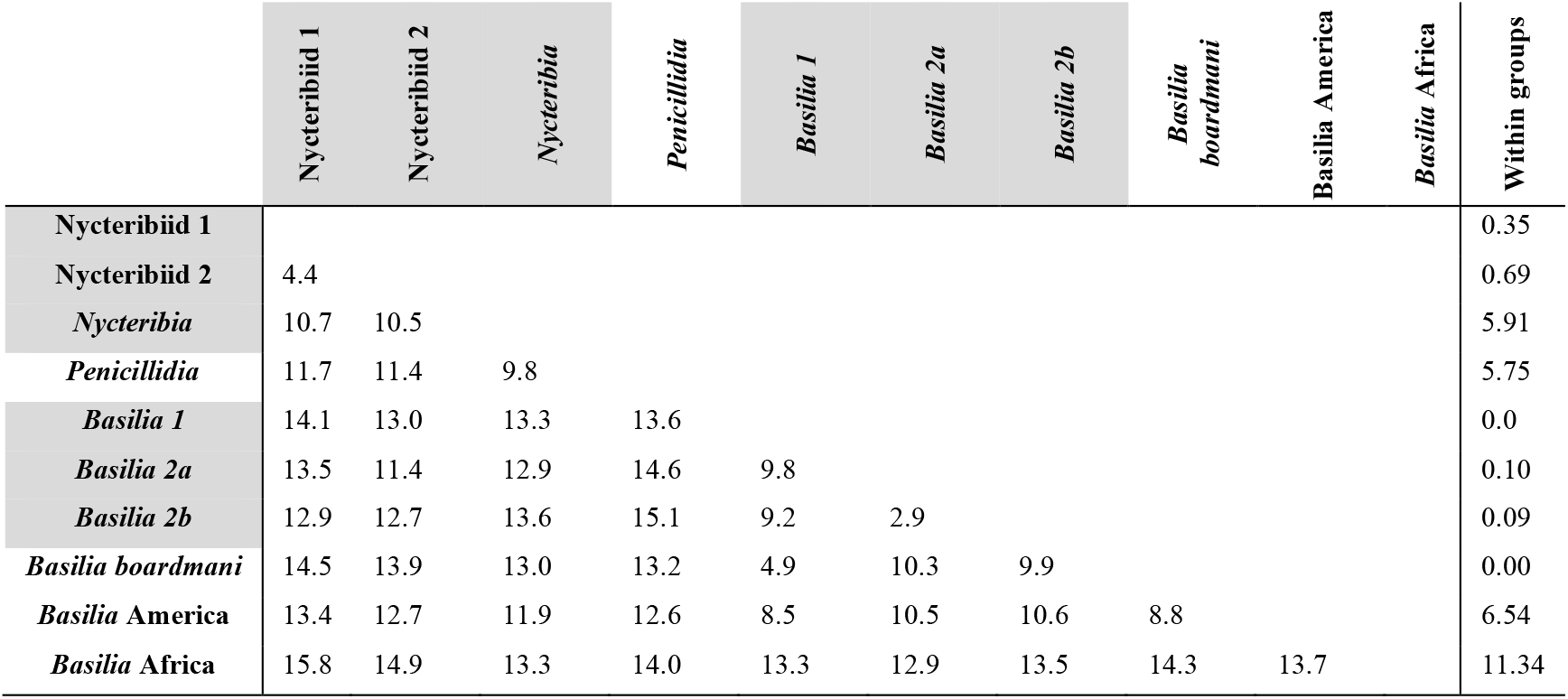
Estimates of evolutionary divergence (COI) in percentage over sequence-pairs between groups (diagonal left matrix), and within groups (most left columns) of bat flies (family Nycteribiidae), showing the lineages obtained in this study (shaded) versus closely related reference sequences from GenBank.

**Appendix 6.**
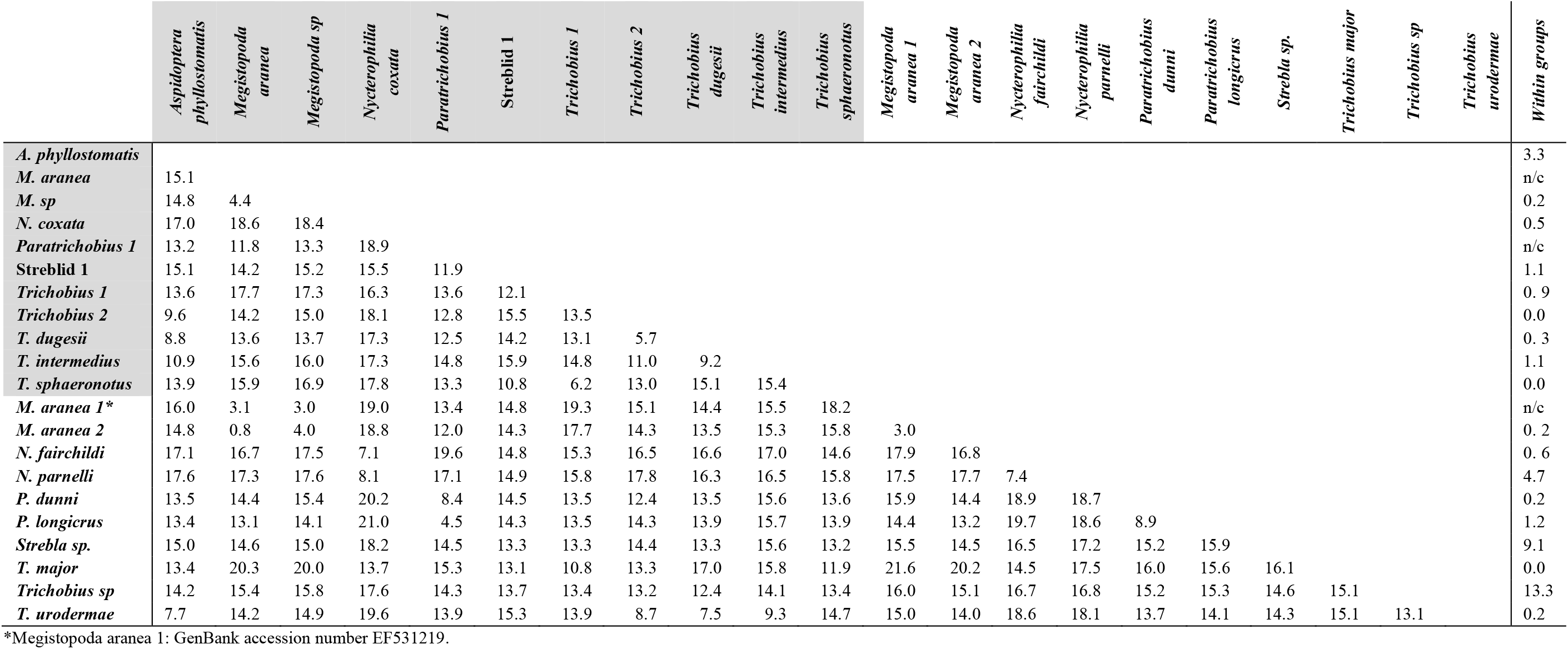
Estimates of percentage divergence (COI) over sequence-pairs between groups (diagonal left matrix), and within groups (most left column) of of bat flies (family Streblidae), showing the lineages obtained in this study (shaded) versus closely related reference sequences from GenBank. Names in the first column have the genus abbreviated.

**Appendix 7.**
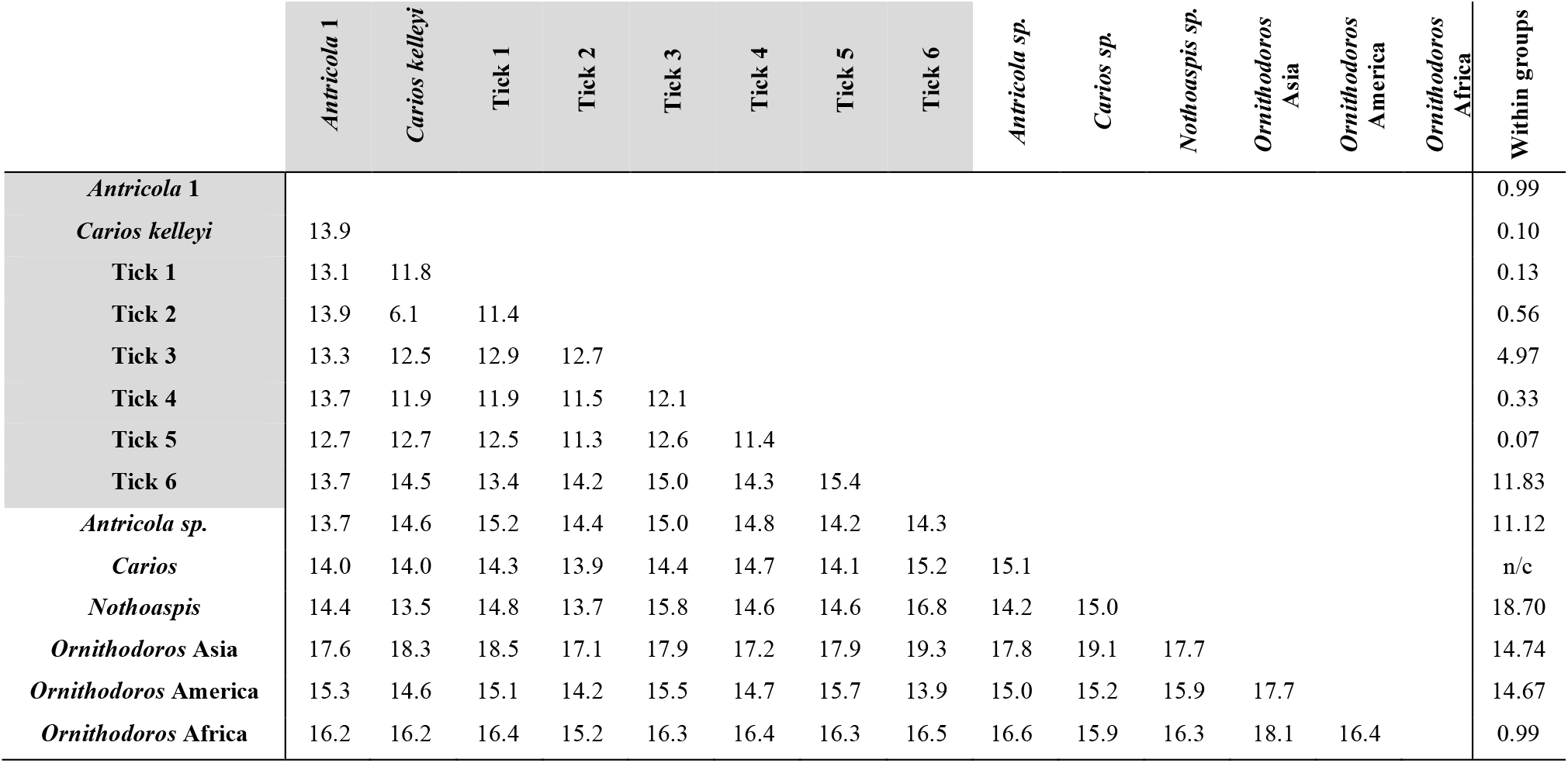
Estimates of evolutionary divergence (COI) in percentage over sequence-pairs between groups (diagonal left matrix), and within groups (most left columns) of bat ticks (family Argasidae), showing the lineages obtained in this study (shaded) versus closely related reference sequences from GenBank.

